# Rv0810c: a genus-conserved, structurally ordered small protein of unknown function carrying DUF3073 in Mycobacterium tuberculosis

**DOI:** 10.64898/2026.08.18.745482

**Authors:** Christophe Guyeux

## Abstract

Small annotated open reading frames are the most neglected part of the functionally uncharacterised *Mycobacterium tuberculosis* genome. We revisit Rv0810c, a 60-residue protein carrying the unknown-function domain DUF3073 (Pfam PF11273), flagged as an Actinobacteria-signature protein in 2006 but never studied since. Rv0810c is genuinely translated (detected in 11 of 16 *Mycobacterium tuberculosis* proteomic datasets), with no significant human homologue, no overlap with a neighbouring gene, and no CRISPR-interference polar effect on either flank. Residue-resolved confidence reveals a bipartite architecture: a rigid 33-residue module (pLDDT 91.9) followed by an extended, acidic, intrinsically disordered tail (radius of gyration 22.8 Å against 11–12 Å expected for a globular protein). The gene is under strong purifying selection (non-synonymous/synonymous ratio 0.86 against 1.93 among 74 size-matched controls, *p* = 5.8 *×* 10^−8^), and its two commonest missense variants are each confined to one sub-lineage, indicating clonal expansion rather than relaxed constraint. DUF3073 is present without a single confirmed loss across 260 well-supported Actinomycetia genera. Despite this conservation, eight independent computational strategies, spanning sequence, structure, electrostatic-patch, embedding-similarity and homo-oligomerisation searches, converge on the same negative: no assignable fold, binding site, or functional neighbour in curated or uncurated sequence space. A phosphosite (Thr24), reproducibly reported by three laboratories, cannot be attributed to a kinase by chemical-genetic or sequence-motif evidence. The contradiction between predicted cytoplasmic topology and reported detection in the macrophage secretory fraction is narrowed, not resolved: ESX secretion, an immunodominant-epitope confound and host-induced transcription are excluded. Rv0810c exemplifies a class of genuinely uncharacterisable small proteins for which negative reporting, not a manufactured function, is the honest outcome.

**Importance:** Many genes of the tuberculosis bacillus still have no known function, and the smallest are the least studied. Rv0810c is one of them: a tiny protein, conserved in essentially every actinobacterium sequenced to date, demonstrably made by the cell, yet without any assignable role after twenty years. We applied eight independent computational approaches and report that all of them fail. Rather than propose a speculative function, we document what is known, what is excluded, and which experiments would settle the question. Publishing such negative dossiers stops other groups from repeating the same dead ends, and identifies the proteins for which laboratory work, not prediction, is now the only way forward.

## 1 Introduction

Whole-genome sequencing of *Mycobacterium tuberculosis* H37Rv in 1998 catalogued nearly 4,000 protein-coding genes, of which several hundred were annotated *hypothetical* or *conserved hypothetical* for want of any functional evidence. A quarter of a century of targeted and systematic follow-up work has substantially reduced that fraction, yet a stubborn residue remains: genes that are short, poorly expressed by the standards of shotgun proteomics, absent from the classical model-organism comparators used for homology transfer, and consequently invisible to nearly every annotation pipeline built around those comparators. Small proteins are disproportionately represented in this residue. Leaderless transcription and translation of very short open reading frames are now recognised as common, not exceptional, features of mycobacterial translation [1], which means that a sizeable population of genuine, translated, potentially essential gene products is systematically under-served by structural and functional genomics resources calibrated on larger proteins.

Rv0810c is a 60-amino-acid protein encoded on the minus strand of the H37Rv chromosome, between *purM* and Rv0811c, and carrying the domain of unknown function DUF3073 (Pfam accession PF11273). The domain was first identified as a signature protein of the phylum Actinobacteria in a comparative-genomics screen of four genomes, including *Mycobacterium tuberculosis* and *Mycobacterium leprae* [2], and confirmed as a broadly distributed actinobacterial marker in subsequent surveys spanning up to 100 genomes [3, 4]. Structural work on a different, unrelated actinobacteria-specific protein of unknown function, ASP1/SCO1997 from *Streptomyces coelicolor*, illustrates both the promise and the limits of this class: an atomic-resolution structure was solved, resembling a nucleoside phosphorylase fold, yet the function remains unknown seventeen years later, because fold resemblance did not translate into a testable functional hypothesis [5]. DUF3073 itself has never been the object of a dedicated functional study in any organism, mycobacterial or otherwise; its only prior mentions in the literature are incidental, as an entry in signature-protein tables [2–4] or as an annotated gene in unrelated genome reports [6].

Two independent lines of experimental evidence give Rv0810c a concrete, if fragmentary, molecular handle that most equally obscure *Mycobacterium tuberculosis* genes lack. First, the protein is phosphorylated at a single threonine, reported by three independent groups using different mass-spectrometry workflows [7–9]. Second, it has been reported in the secretory protein fraction enriched from macrophages infected with *Mycobacterium tuberculosis* [10], an observation seemingly at odds with computational topology prediction, which places the protein in the cytoplasm. Neither observation has previously been pursued beyond the original reporting dataset.

This study asks a single, deliberately narrow question: what, if anything, can be established about Rv0810c using the full range of comparative-genomic, structural, evolutionary and functional computational resources available in 2026, and where does that inventory run out. We treat a documented, honestly reported absence of function as a legitimate and useful outcome, not a failure to be concealed by premature functional or therapeutic framing. Accordingly we deliberately avoid two rhetorical moves common in the dark-proteome literature: presenting a conserved, essential-looking gene as a drug target before its function is established, and inferring essentiality or druggability from measurements that were not designed, or validated, to support such a claim (see Materials and Methods and Discussion for the specific case of a CRISPR-interference vulnerability index that Rv0810c’s own source publication explicitly flags as unreliable for this gene).

## 2 Materials and Methods

### 2.1 Genomic and proteomic resources

Strain-level genomic and epidemiological data (single-position deletion/insertion, SPDI, variant calls; lineage assignments; regions of difference; CRISPR-interference guide coordinates) were drawn from the group’s curated *Mycobacterium tuberculosis* complex (MTBC) database, comprising approximately 255,000 genomes assembled and annotated by a common pipeline (TBannotator) [11], queried through its PostgreSQL and Elasticsearch interfaces at the database snapshot current as of 2026-08. Lineage nomenclature follows the tblearn classification used throughout the database. Proteomic detection evidence was obtained from PaxDb (queried 2026-08) [12], aggregating 16 independent *Mycobacterium tuberculosis* mass-spectrometry datasets. Human homology was assessed by BLASTp (NCBI BLAST 2.17.0+, remote nr database) against the non-redundant protein database restricted to *Homo sapiens* (ENTREZ_QUERY=txid9606[ORGN]), with human insulin (NP_000198.1) as a positive control to confirm that the taxon restriction and search parameters retrieve true positives. Because nr is a living database, the query was run on 2026-08-01 and independently re-run on 2026-08-17 to confirm the negative result was not an artefact of a stale snapshot; the human-restricted sequence count grew from 2,284,585 to 2,284,884 over that interval, consistent with ordinary database growth rather than a search failure.

### 2.2 Structure prediction and residue-resolved confidence analysis

A structural model of Rv0810c (UniProt I6XWB9, AlphaFold Monomer v2.0 preset, entry file dated 2025-08-01) was obtained from the AlphaFold Protein Structure Database [13]. Per-residue predicted local distance difference test (pLDDT) values were extracted and averaged over pre-defined windows (residues 1–19, 20–33, 34–60) to characterise architectural heterogeneity that a single whole-protein average would conceal. Radius of gyration was computed from the predicted coordinates and compared to the expectation for a compact globular protein of 60 residues (11– 12 Å, standard scaling relationship for folded proteins). Remote homology and fold assignment were tested by two independent, non-overlapping routes: profile-profile search with HHsearch 3.3.0 (HH-suite) [14] against the Pfam and SCOPe70_1.75 profile databases, and structure-based search with Foldseek (build 10-941cd33, 3diaa mode, queried 2026-08-10 via the public search.foldseek.com API) against the PDB100, AlphaFold/SwissProt, and CATH50 databases [15], applied both to the full-length protein and to the rigid module (residues 1–33) alone. A third, orthogonal line of evidence was obtained by embedding-based similarity search against the ESM Metagenomic Atlas (queried 2026-08-10), comprising approximately 6.8 billion predicted protein sequences [16], using cosine similarity of sequence embeddings; a within-family positive control (Rv0810c against its *Mycobacterium leprae* orthologue ML2204) and an unrelated-protein negative control were run alongside the query to calibrate the similarity threshold.

### 2.3 Electrostatics, disorder and multiple-sequence-alignment conservation

Charge patterning was quantified by the fraction of charged residues (FCR) and the sequence charge decoration (SCD) parameter, compared to the empirical distribution of 74 size-matched (50–70 residue) H37Rv proteins and to the null expectation obtained by 1,000 intra-sequence permutations of the observed charge sequence. A DUF3073 multiple-sequence alignment of 1,930 curated Pfam PF11273 sequences (retrieved 2026-08) [17] was used to compute per-residue conservation, ranked genome-wide against the 59 alignment columns spanning the full protein, module, hinge and acidic tail alike. Candidate nucleic-acid-binding patches were tested on the AlphaFold model by two independent residue sets, basic residues (K/R) and the six most invariant non-basic module positions, each scored for spatial clustering against a null distribution obtained by randomising residue identity over the folded surface. Intrinsic disorder was additionally flagged by metapredict v3.0.2 [18], cross-checked against the pLDDT profile and radius of gyration described above rather than taken at face value.

### 2.4 Evolutionary constraint and lineage-resolved variant analysis

All missense and synonymous single-nucleotide variants annotated to Rv0810c were retrieved by an exhaustive query of tb_report_spdi_annotations across the full TBannotator strain set (145,209 phenotyped/genotyped strains for allele-frequency denominators), rather than by reusing a hand-curated variant list built for an unrelated purpose. The resulting non-synonymous/synonymous (NS/S) ratio was compared, by two-sided Fisher’s exact test, to the pooled NS/S ratio of the same 74 size-matched H37Rv control proteins used for the charge analysis, and additionally to each control individually. For the two most frequent missense variants, strain-level lineage assignments were cross-tabulated against genome-wide lineage prevalence, and enrichment was tested by Fisher’s exact test with odds ratios, to distinguish a lineage-restricted clonal expansion from a dispersed, recurrent polymorphism. Missense tolerance was independently scored with ESM-1v [19], computing the log-likelihood ratio of each observed substitution and testing for a correlation, by Spearman’s rho, between population frequency and predicted damage.

### 2.5 Genus- and class-wide conservation screen

Presence of DUF3073 was first assessed across 53 non-tuberculous mycobacterial genomes and 13 representative genera outside *Mycobacterium*, using pre-computed orthology and phylostratum layers. To test for genuine gene loss at larger phylogenetic depth, InterPro domain coverage of PF11273 was cross-referenced (NCBI Taxonomy and RefSeq assembly summaries queried 2026-08) against all genera of the class Actinomycetia with at least one reference-quality RefSeq assembly (260 of 491 candidate genera after excluding taxonomically divergent classes erroneously included under a broad *Actinobacteria* NCBI subtree query: Coriobacteriia, Rubrobacteria, Acidimicrobiia and Thermoleophilia, none of which carry the domain). Genera flagged as apparently domain-negative by this screen, and supported by at least two independent assemblies, were re-examined by direct BLASTp against their proteomes to distinguish a true loss from a detection-threshold false negative.

### 2.6 CRISPR interference, homo-oligomerisation and functional-context data

CRISPR-interference knockdown data were taken from the genome-wide dataset of Bosch et al. [20], using the raw per-guide depletion measurements (23 guides targeting Rv0810c, entirely contained within the gene and unique in the genome) rather than the derived vulnerability index (VI), which the source publication itself classifies as unreliable (certain = False) for this gene owing to a below-median guide count and a narrow range of achieved knockdown strengths; we report guide-level depletion and flatline rate instead, and discuss this methodological distinction explicitly (Discussion). A published dose-response re-analysis of the Bosch screen [21] was additionally consulted as a counter-argument check. Homo-oligomerisation was tested by Boltz-2 (v2.2.1) structure prediction of the Rv0810c homodimer (five independent models), with the ribosomal protein RpmG2/Rv0634B, known not to self-associate outside the assembled ribosome, as a positive-artefact control for the pipeline’s propensity to report spuriously confident interfaces. Transcript-level regulatory context (a candidate 5*^′^* untranslated region/leader element, ncRv10810c) was assessed for cross-species DNA conservation and for thermodynamically stabilised secondary structure relative to a randomised sequence null model. Genome-wide, condition-resolved transcriptional behaviour was assessed using the independent-component-analysis compendium of Yoo et al. [22], downloaded from the Reosu/modulome_mtb GitHub repository in 2026-08 (647 RNA-seq profiles across 231 conditions, including THP-1 and alveolar macrophage infection, murine bone-marrow-derived macrophages, hypoxia, dormancy, resuscitation, nitrate and redox stress, sputum, lipid-body and biofilm states), comparing the mean expression of Rv0810c across 125 host-associated samples to 27 standard-growth samples, and ranking the resulting delta against the same 74-protein size-matched control set.

### 2.7 Localisation contradiction: ESX secretion motif and immune-epitope screening

Subcellular localisation was predicted with DeepTMHMM [23] (BioLib-hosted model, queried 2026-08), to test for a signal peptide or transmembrane segment consistent with extracellular or membrane targeting. A candidate general ESX/type-VII secretion signal (Y-x-x-x-[D/E], typically located in the unstructured C-terminal segment of WXG100 substrates or their dimer partner) [24] was scanned across the full length of Rv0810c and, as positive and negative controls, across three confirmed ESX-1 substrates retrieved from UniProt (EsxA/ESAT-6, P9WNK7; EsxB/CFP-10, P9WNK5; EspB, P9WJD9). Candidate immunodominant epitopes were queried directly (2026-08) from the Immune Epitope Database (IEDB) application programming interface (query-api.iedb.org), filtering B-cell and T-cell assay records by parent source antigen UniProt accession; every returned peptide was re-localised on the H37Rv reference sequence rather than trusted at its reported coordinate, following a data-integrity issue encountered repeatedly in this project’s other supplementary-table sources (see Discussion).

The bioinformatic analyses, database queries and literature cross-referencing underlying this study were carried out within an AI-augmented bioinformatics working environment, Claude Code (Anthropic), operating the Claude Opus 5 language model as an interactive analysis agent under continuous human direction. Every script, statistical test and numerical claim produced within this environment was independently traced back to its underlying raw data, primary publication or database record before being reported here, following the verification discipline described throughout this section; the corresponding author designed the study, specified and reviewed every analysis, and takes full responsibility for the accuracy of its content.

## 3 Results

### 3.1 Rv0810c is a bona fide, translated, human-non-homologous protein with no confounding genomic overlap

Rv0810c is detected in 11 of the 16 independent proteomic datasets aggregated by PaxDb, indicating that the gene is translated under standard laboratory growth conditions rather than being a purely computational annotation artefact. BLASTp against the human-restricted non-redundant protein database returned no significant hit against any of 2,284,585 human sequences, while the positive control (human insulin) returned the expected matches under identical search parameters, confirming that the negative result reflects true absence of homology rather than a failed search. The gene shows no nucleotide overlap with either flanking gene (0%), and all 23 CRISPR-interference guides used in the genome-wide screen described below are uniquely mapped and entirely contained within the Rv0810c coding sequence, ruling out a polar knockdown effect on Rv0811c or Rv0812 as a confound for any phenotype attributed to Rv0810c.

### 3.2 A bipartite architecture: a rigid module and an acidic disordered tail

The AlphaFold model reports a whole-protein mean pLDDT of 77.0, a value that would conventionally be read as a moderately confident, largely folded prediction. Residue-resolved analysis shows this average to be an artefact of pooling two structurally distinct halves: residues 1–19 average pLDDT 91.9 (a high-confidence, rigid module, invariant across the phylum), residues 20–33 average 85.6, and residues 34–60, the tail, average only 62.0. The predicted radius of gyration is 22.8 Å, roughly double the 11–12 Å expected for a compact globular protein of 60 residues, indicating that the molecule as a whole is extended rather than folded into a single compact domain. This bipartite reading is corroborated by an independent line of evidence: intrinsic-disorder prediction by metapredict flags disorder broadly across the sequence, which taken alone might be dismissed as a false positive given the pLDDT of the rigid module, but is in fact consistent once restricted to the tail, where three independent signals converge, low pLDDT, a positive metapredict call, and marked hypervariability of tail length across species (from approximately 8 residues in *Tropheryma whipplei* to approximately 50 in *Streptomyces coelicolor* ). This architecture parsimoniously explains three otherwise separate negative results reported below: the absence of a localised electrostatic binding patch, the absence of confident homo-oligomerisation, and the failure of embedding-based functional similarity search, all of which implicitly assume, or are best powered to detect, a single compact folded surface.

**Figure 1:**
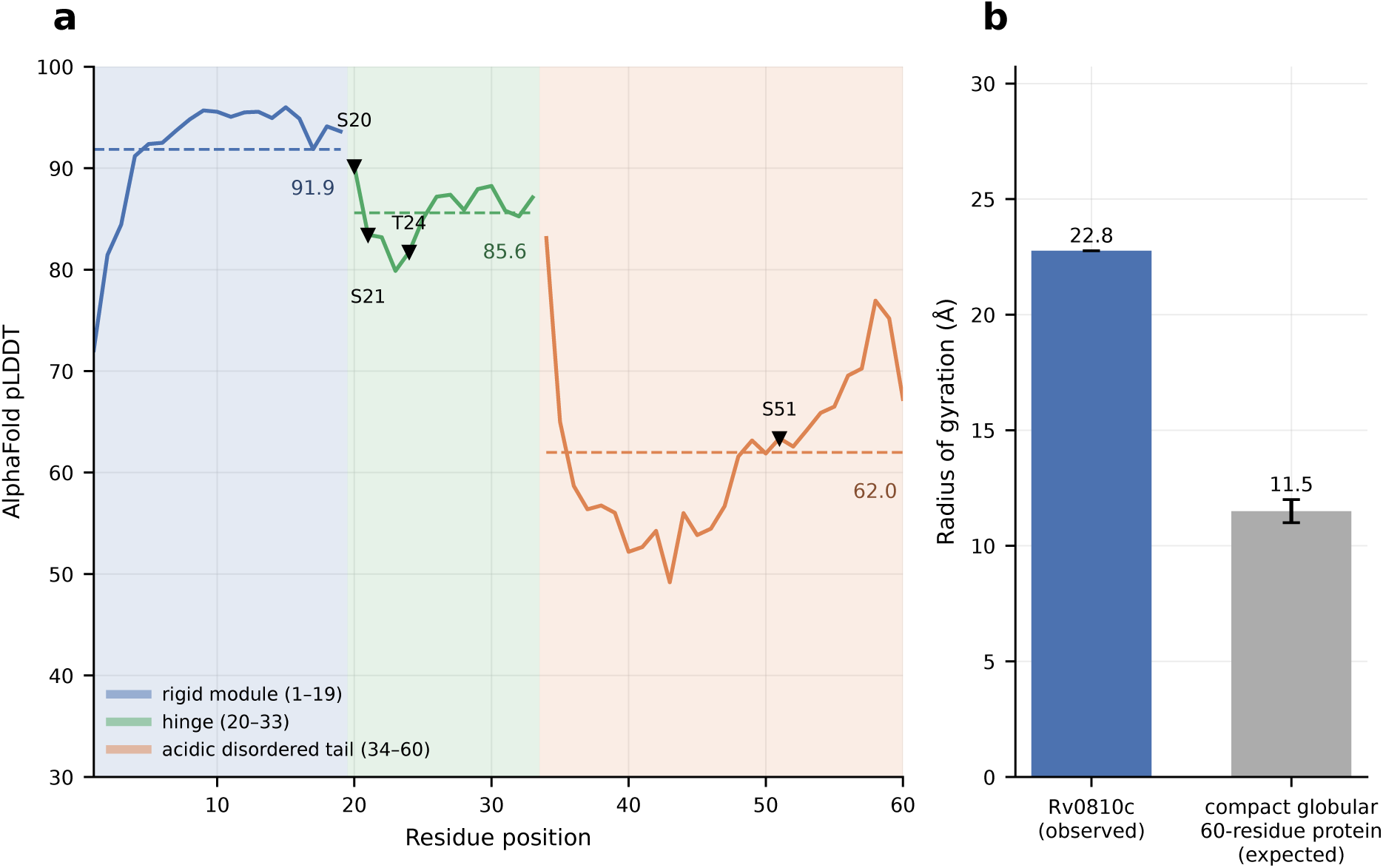
A bipartite architecture separates a rigid module from an acidic disordered tail. (**a**) Per-residue AlphaFold pLDDT profile, coloured by the three windows analysed (rigid module, residues 1–19; hinge, 20–33; acidic disordered tail, 34–60), with the mean pLDDT of each window shown as a dashed line. Triangles mark the four residues discussed as phosphorylation sites (Section 3.6): Ser20, Ser21 and Thr24 fall in the hinge, Ser51 in the tail. (**b**) Radius of gyration computed from the predicted C*α* coordinates, compared to the expectation for a compact globular protein of 60 residues (11–12 Å, error bar spans this range); the roughly two-fold excess indicates an extended, non-globular molecule rather than a single compact domain.

The acidic tail itself is under compositional, not length, constraint: its fraction of aspartate and glutamate residues is independent of tail length across species (Spearman *ρ* = 0.09, not significant; median fraction 35.4%), while the length itself varies by close to an order of magnitude across clades without correlation to host genome size (*ρ* = *−*0.08 across 16 taxonomic orders). No stable intramolecular electrostatic interaction was detected between the rigid module and the tail (minimum inter-domain C*α* distance 29.0 Å, no contacts below 12 Å), ruling out a simple head-to-tail electrostatic clamp as the functional rationale for this bipartite design.

Charge patterning within the rigid module is itself atypical for a protein of this size: the fraction of charged residues (FCR = 0.417) ranks second among 75 H37Rv proteins of 50–70 residues, despite a near-neutral net charge (rank 71/75), indicating a polyampholyte rather than a net-charged polypeptide. Charge is significantly more clustered along the sequence than expected from its own composition (sequence charge decoration = *−*3.18 against a permutation null of *−*0.72, *p* = 0.0022, *z* = *−*5.24), a patterning signature more often associated with disorder-driven electrostatic function than with a rigid folded module, and one that the present study was unable to connect to any detectable binding activity (Section 3.5).

### 3.3 DUF3073 conservation marks a structural boundary, not a simple avoidance of basic residues

Among the 1,930 sequences of the curated DUF3073 (PF11273) alignment, seven positions of the rigid module are perfectly invariant, zero observed substitutions among aligned sequences (majority-residue frequency 100%, alignment-column ranks 1–7 of 59, tied and separated here only by residue order): G2, R3, G4, R5, A8, A14 and L29. Two of these seven, R3 and R5, are basic; the module’s third basic residue, K18, is very slightly but measurably less conserved (99.95% majority frequency, rank 8/59), as are two further module lysines, K9 (99.89%, rank 10/59) and K7 (99.37%, rank 11/59). E32, immediately adjacent to the order-to-disorder boundary described below, is conserved at 99.0% (rank 12/59). Basic residues are therefore not uniformly under weaker sequence constraint than the module’s other invariant positions: two of its three most conserved basic residues are exactly as invariant as its most conserved non-basic positions, and only K18, K9 and K7 show any measurable, if slight, relaxation. This tempers rather than supports a simple electrostatic-avoidance reading of the module’s conservation pattern; whatever functional constraint dominates this domain’s evolution is not reducible to a uniform pressure against basic residues. E32 and the adjacent L33 do mark, at the level of sequence conservation, the same order-to-disorder boundary independently identified structurally by the pLDDT and radius-of-gyration analysis above, a convergence between two entirely independent methods (structure prediction and multiple-sequence-alignment statistics) that we consider one of the more robust architectural conclusions of this study.

### 3.4 Rv0810c is under strong, statistically formalised purifying selection, and its two commonest missense variants are traceable clonal expansions

An exhaustive query of the full TBannotator variant catalogue identifies 84 distinct missense sites in Rv0810c, including two above a 0.01% population-frequency threshold: p.Asp56Ala (73/145,209 strains, 0.050%) and p.Thr24Pro (32/145,209 strains, 0.022%). The resulting non-synonymous/ synonymous ratio (86 non-synonymous versus 100 synonymous sites, NS/S = 0.86) is markedly lower than the pooled ratio of 74 size-matched H37Rv control proteins (NS/S = 1.93), placing Rv0810c at the 2.7th percentile of the control distribution, the second most non-synonymous-depleted protein of the entire comparator set (Fisher’s exact test against the pool, *p* = 5.8 *×* 10^−8^; 65 of 74 individual pairwise comparisons independently significant in the same direction, none in the opposite direction).

Both frequent missense variants are strongly lineage-restricted rather than dispersed across the phylogeny. p.Asp56Ala is 94.5% concentrated in sub-lineage L4.1.1.1 (which itself represents only 1.26% of the TBannotator strain collection; Fisher’s exact odds ratio = 1387, *p* = 3.3 *×* 10^−126^), yet only 2.21% of L4.1.1.1 strains actually carry the variant, ruling out fixation within the sub-lineage. p.Thr24Pro is 96.7% concentrated in sub-lineage L4.3.3 (5.81% of the collection; odds ratio = 471, *p* = 3.9 *×* 10^−35^), carried by only 0.20% of L4.3.3 strains. In both cases, the single most heavily sampled lineage in the entire database, L2.2.1 (71,496 strains), harbours almost no carriers (zero and one strains respectively), which directly rules out a global sampling-depth artefact as an alternative explanation for the observed concentration. This pattern is consistent with each variant having arisen once (or a small number of times) and expanded clonally within a restricted transmission chain, rather than reflecting recurrent, functionally tolerated substitution; a formal distinction between a single founder event and localised relaxation of selection would require pairwise transmission-distance analysis among carriers, which was not performed here. ESM-1v scoring of the full 84-site missense catalogue and of a saturation scan of the rigid module finds no correlation between observed population frequency and predicted mutational damage (whole protein *ρ* = *−*0.111, *p* = 0.32; module *ρ* = *−*0.084, *p* = 0.65; tail *ρ* = *−*0.096, *p* = 0.51), and substitutions at the three characterised phosphorylatable residues (Ser20, Ser21, Thr24; Section 3.6) are not more damaging, by this measure, than the module average (permutation test, *p* = 0.97, one-sided). This does not contradict the functional relevance of Thr24 phosphorylation: ESM-1v scores tolerance across all 19 possible substitutions, not the narrower constraint of remaining a phosphorylatable serine or threonine that a genuine phosphosite would be expected to impose, and the two measures are simply addressing different questions.

### 3.5 DUF3073 is present across the entire well-sequenced class Actinomycetia, without a single confirmed loss

Rv0810c orthologues were detected in all 53 tested non-tuberculous mycobacterial genomes and in 10 of 13 tested genera outside *Mycobacterium*; the one apparent exception, *Streptomyces coelicolor*, proved to be a tblastn detection-threshold false negative, resolved by direct Pfam/ InterPro annotation (UniProt Q9RKJ8). Extending the screen to the entire class Actinomycetia (260 of 491 candidate NCBI genera retained after excluding four taxonomically divergent classes erroneously captured by a broad *Actinobacteria* subtree query, none of which in fact carry the domain), InterPro coverage of PF11273 flagged 17 apparently domain-negative genera, of which four are supported by two or more independent reference-quality assemblies (*Solwaraspora*, *Spirillospora*, *Carbonicoccus*, *Oryzobacter* ). Direct BLASTp against these four genera’s proteomes confirmed, in all four cases, that the gene is in fact present and already annotated as a “DUF3073 domain-containing protein” by RefSeq/PGAP (e-values down to 4 *×* 10^−15^), meaning that the InterPro screen itself, not the underlying biology, produced the apparent gaps. No genuine loss of DUF3073 was identified anywhere in the 260 well-supported genera examined; the remaining 13 single-assembly candidates were not pursued further, as the yield from the four best-supported candidates was uniformly zero. DUF3073 can therefore be described, on the evidence assembled here, as universal across the class Actinomycetia to the resolution of currently available reference-quality genome assemblies.

### 3.6 Eight independent computational approaches converge on the same negative: no assignable molecular function

No fold could be assigned to Rv0810c, or to its rigid module in isolation, by either of two independent structure-comparison strategies. Profile-profile search (HHpred) against the rigid module alone returned a best non-self match of 40.5% probability against Pfam and 25.4% against SCOPe70, both far below conventional confidence thresholds for a genuine homology call. Structure-based search (Foldseek) against three reference databases (PDB100, AlphaFold/SwissProt, CATH50) returned zero hits for the module in isolation. A third, orthogonal approach, embedding-based similarity search against the ESM Metagenomic Atlas (6.8 billion sequences), returned a strong positive-control signal (cosine similarity 0.957 between Rv0810c and its *M. leprae* orthologue ML2204, against 0.211 for an unrelated protein), confirming that the search itself is capable of detecting true relationships at this scale, yet none of the 25 nearest neighbours of Rv0810c in this vast, uncurated sequence space corresponds to an annotated protein; all are raw metagenomic entries. DUF3073 is therefore not merely an orphan family with respect to curated structural and functional databases; it appears to be an orphan family with respect to the entire publicly available protein universe, curated or not.

No localised nucleic-acid-binding surface could be detected on the rigid module by any of four independent tests: a sequence-motif scan (ScanProsite, zero hits), profile-profile search (nothing interpretable), and two structural clustering tests, one restricted to basic residues (*z* = 0.06, *p* = 0.49, no patch) and one restricted to the six most invariant non-basic module positions identified in Section 3.3 (*z* = +1.53, *p* = 0.951, more spatially dispersed than a random null, not less). A non-localised, distributed binding mode along the extended structure cannot be excluded by these tests and would require wet-laboratory validation (electrophoretic mobility shift assay, pulldown) that falls outside the scope of this computational study.

Controlled homo-oligomerisation prediction with Boltz-2 found no evidence for a Rv0810c homodimer (interface predicted TM-score, ipTM, 0.091–0.138; inter-chain predicted aligned error 15.9–17.0 Å; complex pLDDT 0.46–0.49; consistent across five independent models), scores well below any conventional confidence threshold. Critically, these scores are lower than those obtained for a positive-artefact control run through the identical pipeline: the ribosomal protein RpmG2/Rv0634B, which is not known to self-associate outside the assembled ribosome, nonetheless returns a confident interface (ipTM 0.385–0.714, predicted aligned error 3.4–8.8 Å, complex pLDDT 0.72–0.77) that this project has previously documented as a generic scoring artefact of the same pipeline on an unrelated protein. That Rv0810c scores below a known false-positive control, rather than merely below an arbitrary threshold, rules out the alternative explanation that the negative result reflects an instrument unable to report confident dimers in this size and abundance regime.

Together, these results extend a previously reported five-method negative inventory (sequence motif, profile-profile, structure search, electrostatic patch/contiguity, controlled homooligomerisation) to eight independent computational strategies, all negative, across both curated and uncurated sequence and structure space.

### 3.7 A reproducibly reported phosphosite of unassignable kinase, in an otherwise nearly closed regulatory context

A single phosphorylated residue, Thr24, is reported across four independent datasets from three laboratories using distinct experimental workflows [7–9], and is conserved without a single gap in 81.3% of the 1,930 PF11273 alignment sequences (Figure 2), a degree of cross-species conservation well above what is typically expected of an incidental, non-functional phosphorylation event.

**Figure 2:**
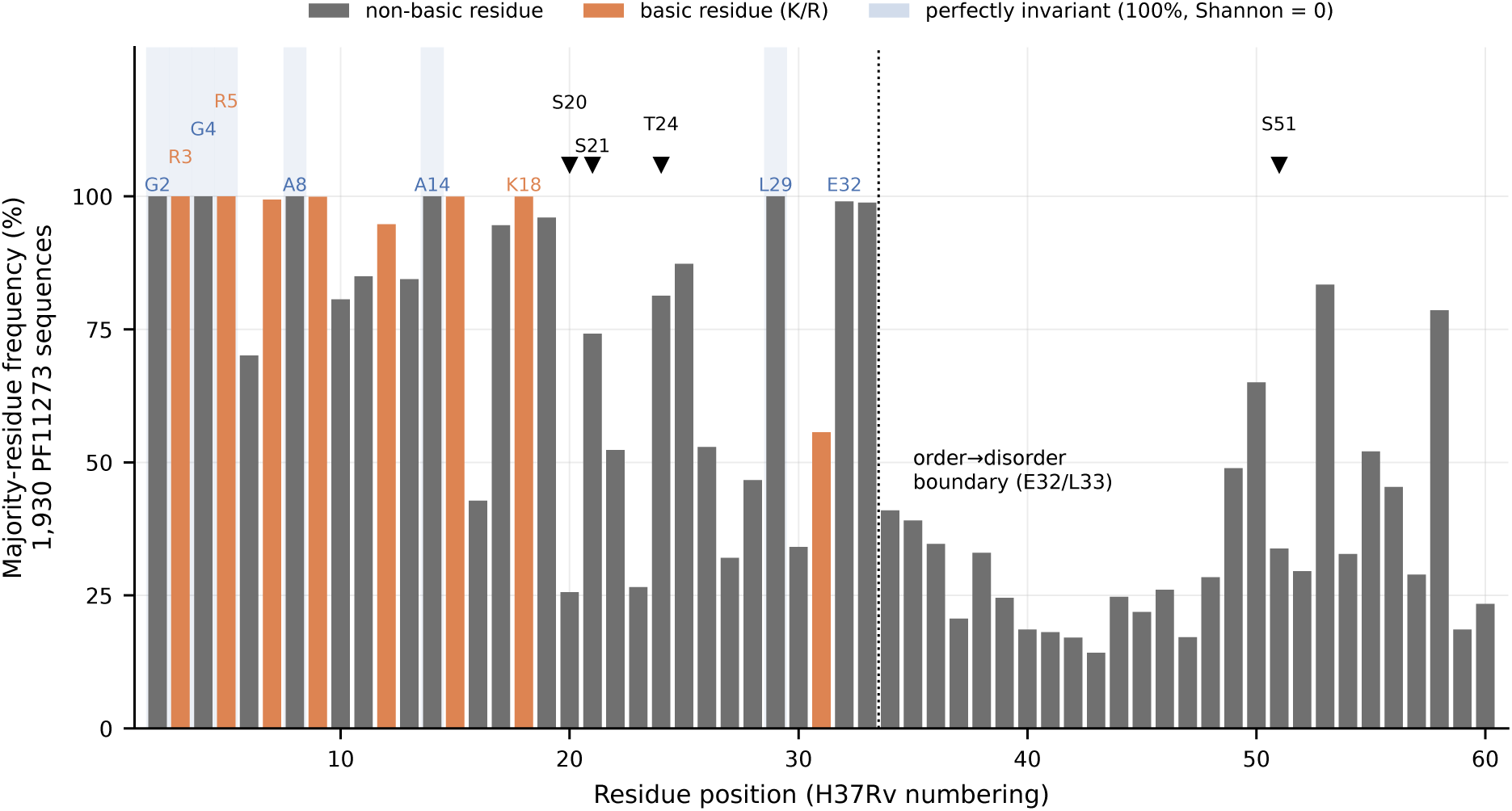
Conservation is deepest at the order-to-disorder boundary, and does not track basic versus non-basic residue identity. Majority-residue frequency at each of the 59 PF11273 alignment columns spanning the full protein, module, hinge and acidic tail alike (1,930 sequences), coloured by whether the H37Rv residue at that position is basic (orange, K/R) or not (grey). Shaded columns mark the seven positions at exactly 100% majority frequency (zero Shannon entropy): two of the module’s three basic residues, R3 and R5, are perfectly invariant alongside the five non-basic positions previously highlighted (G2, G4, A8, A14, L29); only K18 among the module’s basic residues falls measurably, if slightly, below this ceiling. The dotted line marks the order-to-disorder boundary after L33 independently identified by the structural analysis of Figure 1. Triangles mark the four phosphorylation sites discussed in Section 3.6.

Two earlier, independent mycobacterial phosphoproteomic surveys [25, 26] did not detect this or any other Rv0810c phosphosite, a pattern more consistent with condition- or strain-dependent detection sensitivity than with a spurious call shared by the three positive datasets. Combined pharmacological inhibition of the serine/threonine kinases PknA and PknB reproducibly and directionally reduces Thr24 phosphorylation in an independent chemical-genetic dataset [8], but the inhibitor used is not selective for a single kinase, so this result cannot attribute the modification to either kinase individually. A second, entirely independent test based on sequence-motif matching against the published consensus of the six characterised *Mycobacterium tuberculosis* serine/threonine protein kinases [26] finds that the local sequence context of Thr24 (YSSPQTDFQR) is a poor match to that consensus, failing the characteristic acidic run at positions *−*4 to *−*1 (0 of 4 acidic residues) and lacking the expected bulky hydrophobic residue at position +3; internal controls (two neighbouring serines, S20 and S21, sharing the same local context) fail the same test in the same way, while an unrelated tail serine (S51) passes by simple compositional coincidence, confirming that the test discriminates meaningfully rather than failing uninformatively. Prisic and colleagues themselves note that the shared consensus motif of the six kinases they characterised “limits direct mapping of individual substrates to a cognate kinase” [26], a limitation that this analysis independently reproduces for Thr24 specifically. Two independent lines of evidence therefore converge on the same conclusion: the kinase responsible for Thr24 phosphorylation cannot currently be assigned with the publicly available resources, a result we read as informative rather than as a simple gap in effort.

A candidate transcribed element in the 5*^′^* intergenic region, 27 nucleotides upstream of the start codon and carrying a detectable *−*10 promoter-like motif, was independently identified at the same coordinates (to within one nucleotide) by two genome-wide *Mycobacterium tuberculosis* transcriptomic surveys [27, 28], expressed in 4 of 15 tested conditions [28]. It is conserved at 92.6% nucleotide identity in *Mycobacterium marinum*, but shows no evidence of a thermodynamically selected secondary structure (*z* = *−*1.15 against a randomised sequence null), favouring a leader/5*^′^* untranslated-region reading over a discrete *trans*-acting small regulatory RNA; it is also absent from the 24 experimentally validated small RNAs catalogued by Ami et al. [28]. Neither reading has been confirmed experimentally (5*^′^* rapid amplification of cDNA ends or Northern blotting). We note, and explicitly do not repeat, a previously published claim that the neighbouring intergenic region between *purM* and Rv0810c itself expresses a small RNA [29]: that statement carries no supporting citation in its source publication, and the same publication’s own supplementary table of small-RNA-expressing intergenic regions does not include this locus, indicating the claim should not be propagated. This does not mean the *purM* -Rv0810c intergenic region is transcriptionally silent: an unnamed element is independently called there by one of the two surveys above (94 nt, antisense to the 3*^′^* end of *purM*, overlapping the Rv0810c 3*^′^*UTR, adjusted *p* = 2.8 *×* 10^−3^) [27], but it is a different element, in the opposite orientation, from the one Zeng and colleagues describe, and it is absent from the second survey [28].

Genome-wide transcriptomic profiling across 231 experimental conditions finds Rv0810c neither induced nor repressed under host-associated conditions relative to standard growth (delta = +0.28 log-TPM between 125 host-associated and 27 standard-growth samples, 66.7th percentile of the same 74-protein size-matched control distribution, which ranges from *−*1.48 to +1.57 with a median close to zero). A gene under this degree of constitutive purifying selection therefore appears to be constitutively expressed rather than transcriptionally responsive to infection, consistent with a housekeeping-like role rather than a dedicated host-interaction function, though this remains an inference from expression stability rather than a direct functional demonstration.

### 3.8 The cytoplasmic-prediction/macrophage-detection contradiction narrows by elimination, not by positive proof

Rv0810c is computationally predicted to be cytoplasmic (no signal peptide or transmembrane segment detected by DeepTMHMM [23]), yet has been reported both as phosphorylated in whole-cell and culture-filtrate fractions [9] and as enriched in the secretory protein fraction of infected macrophages [10]. Three concrete, testable mechanisms that could reconcile a cytoplasmic protein with extracellular or macrophage-fraction detection were examined and, in each case, ruled out. First, a scan for the general ESX/type-VII secretion signal [24] found no match anywhere across the 60 residues of Rv0810c, in either the C-terminal window where the signal is functionally located in true substrates or elsewhere; the same scan correctly identified the functional signal in the positive control (EsxB/CFP-10, within its expected C-terminal window) and distinguished it from a positionally irrelevant, fortuitous match in a second control (EspB), showing that the test itself is discriminating rather than uninformative. This negative result is consistent with the earlier structural finding that Foldseek returns zero hits for Rv0810c against a database that includes the WXG100-fold ESX substrates, and with the absence of any physical genomic linkage between Rv0810c and an ESX locus. Second, a direct query of the Immune Epitope Database identified only two relevant references covering 30 of the protein’s 60 residues (two peptides, spanning positions 25–39 and 45–59, the latter overlapping the p.Asp56Ala missense hotspot of Section 3.4); every returned T-cell record (four, after deduplication and re-localisation on the reference sequence) was scored “Negative” for HLA class II reactivity, including in a large contemporary dataset [30], providing no support for a documented immunodominant-epitope confound on the fraction tested, though Thr24 and the entire N-terminal module (residues 1–24) were never covered by any assay in this database and remain untested. Third, the transcriptomic compendium analysis of Section 3.6 found no evidence of host-induced transcriptional up-regulation that might otherwise explain increased protein abundance during infection.

None of these three eliminations provides a positive mechanistic replacement for the observed macrophage-fraction detection. The most parsimonious remaining reading, that a macrophage secretoryfraction preparation is contaminated by lysis of a small, abundant cytoplasmic protein, a well-known limitation of such fractionation protocols, is reinforced by the elimination of three concrete alternatives, but it is not directly demonstrated, and the contradiction should be considered narrowed rather than resolved.

## 4 Discussion

Taken together, these results describe a small protein that is unambiguously real, deeply conserved, architecturally distinctive, and under some of the strongest sequence-level purifying selection observed among comparably sized *Mycobacterium tuberculosis* proteins, while remaining functionally opaque to every computational strategy applied to it. We consider this combination scientifically informative rather than a null result to be minimised. Two aspects of the dossier deserve particular emphasis.

First, the negative functional inventory is now broad enough, eight independent methods spanning sequence motif analysis, two orthogonal profile-profile and structure-search strategies, embedding-based similarity search across the largest publicly available uncurated sequence space, electrostatic patch detection restricted to two independently justified residue sets, and a controlled homooligomerisation test validated against a known artefact-producing positive control, that continued undirected computational search is unlikely to be productive. Any future progress on the molecular function of DUF3073 will most plausibly come from wet-laboratory approaches (targeted pulldown, crosslinking mass spectrometry, or structural determination of the full-length protein including its disordered tail, which AlphaFold-type single-chain prediction is inherently poorly suited to resolve) rather than from further *in silico* screening. This mirrors the precedent set by ASP1/SCO1997, a different actinobacteria-specific protein of unknown function for which an experimentally determined structure was obtained seventeen years ago without yielding a functional assignment [5]: structural determination narrows but does not by itself resolve function for this class of protein, and DUF3073, which additionally lacks any assignable fold at all, sits at a still earlier stage of that trajectory.

Second, we deliberately do not frame Rv0810c as a drug target, and we consider this restraint methodologically important rather than merely cautious. The strongest quantitative signal available for this gene under CRISPR interference, its published vulnerability index, is explicitly flagged as unreliable for Rv0810c by the index’s own source publication (certain = False), a consequence of a below-median guide count (23 against a median of 74 for confidently essential genes) concentrated in a narrow range of achieved knockdown strengths, which propagates into unusually wide parameter confidence intervals. What remains defensible from the same dataset is considerably more modest: the 23 guides targeting Rv0810c behave normally by the screen’s own internal quality metrics (near-zero baseline read-count skew, a genuine mean depletion effect, and a flatline rate matching that of a size-matched comparator panel), indicating that the gene is depleted upon knockdown, without that depletion being reducible to a single, citable vulnerability estimate. An independent published re-analysis of the same screening data [21] in fact singles out Rv0810c as an example of a low-abundance, high-noise gene, significant under one statistical model but not under a more conservative dose-response model, a nuance that should temper, not resolve, the essentiality question; low baseline transcript or guide-representation abundance is, after all, also the expected consequence of strong vulnerability, so the two readings are not straightforwardly reconcilable from the published data alone. Direct transposon-insertion evidence is similarly thin, five TA dinucleotide sites, four of them permissive, a limitation shared by 96% of *Mycobacterium tuberculosis* genes in the 150–300 nucleotide size range and therefore not itself anomalous, but not a basis for a strong essentiality claim either. The orthologous gene in *Corynebacterium diphtheriae* is independently reported essential by transposon-directed insertion sequencing and mass spectrometry [31], a genuinely independent line of evidence from a different organism and methodology, and the one piece of the essentiality case we consider reasonably solid; sequence-level purifying selection, by contrast, is now solidly established by the present study (*p* = 5.8 *×* 10^−8^), but constraint and essentiality are conceptually distinct properties, and we have kept that distinction explicit throughout rather than allowing the stronger of the two claims to stand in for the weaker.

A further published association, a genome-wide association study hit linking Rv0810c to rifampicin and rifabutin resistance [32], does not survive a population-structure control applied here for the first time to this locus: the only drug association independently supported in the original variant-calling study for this intergenic region is with ethionamide, not rifampicin [29], and a direct query of the TBannotator database finds zero rifampicin-resistant strains among 59 phenotyped carriers of the ten most frequent variants in this region (Fisher’s exact test, *p* = 1.0, odds ratio = 0), with the two most discriminating variants each corresponding to a pure lineage marker. This is consistent with genetic hitchhiking driven by population structure rather than a causal drug-resistance mechanism, and illustrates a recurring pattern in this dossier: several of the strongest-sounding published associations for this locus, an intergenic small RNA, a resistance GWAS hit, a curated phosphosite position, dissolve or require substantial correction on direct re-examination of primary source data or tables, a caution we would extend to any future literature-based claim about this gene.

The remaining open question, the mechanism linking Thr24 phosphorylation and reported macrophage-fraction detection to an otherwise cytoplasmic protein, illustrates the limits of computational elimination as a method: three concrete, testable hypotheses were ruled out, yet no positive replacement mechanism was identified, leaving the simplest explanation, fractionation artefact from lysis of an abundant cytoplasmic protein, as the best-supported but not proven reading. We consider this an appropriate place for the computational component of this investigation to stop, and an appropriate handoff point to dedicated wet-laboratory work (live-macrophage proteomics, immunoprecipitation) rather than further *in silico* inference.

## 5 Conclusion

Rv0810c is a genuine, translated, deeply conserved small protein carrying a domain of unknown function, DUF3073, that is universal across the entire class Actinomycetia to the resolution of currently available reference genomes, yet has never been functionally characterised in any species in twenty years since its first description as a signature protein of this phylum. An extensive, methodologically diverse computational investigation, spanning sequence, structure, evolutionary constraint, regulatory context and cross-species conservation, converges on a single conclusion: no molecular function can currently be assigned to this protein by any available *in silico* approach, and its strongest quantitative property, strong purifying selection, characterises the gene as biologically important without indicating what it does. We report this outcome in full rather than manufacturing a functional narrative from suggestive but unresolved fragments (a phosphosite of unidentified kinase, a localisation contradiction narrowed but not resolved, a CRISPR-interference signal that its own source data classify as not reliably quantifiable). We consider Rv0810c a well-documented, tractable starting point for future experimental work on DUF3073, and more broadly an illustration that a rigorously negative computational dossier, honestly reported, is itself a useful and publishable contribution to closing the dark proteome of *Mycobacterium tuberculosis*.

## Data, Metadata, and Code Availability

All data analysed in this study are publicly available and are cited at each use throughout the Materials and Methods, with accession numbers, DOIs and supplementary-data sources given in place. The principal sources are the H37Rv reference annotation (NC_000962.3), the curated DUF3073 alignment of Pfam PF11273, AlphaFold DB and UniProt records, InterPro and NCBI RefSeq assembly summaries, PaxDb and the published *Mycobacterium tuberculosis* proteomic datasets cited in the text, the CRISPR-interference library of Bosch et al. [20], the Immune Epitope Database, and the transcriptomic compendium of Yoo et al. [22]. Population variation was queried from a curated MTBC variant database. Analysis scripts and the derived data required to regenerate every figure and every reported statistic are deposited at https://github.com/cguyeux/deciphering-tuberculosis-with-ai and archived at https://doi.org/10.5281/zenodo.21988886.

## CRediT authorship contribution statement

Christophe Guyeux: Conceptualization, Data curation, Formal analysis, Investigation, Methodology, Project administration, Resources, Software, Validation, Visualization, Writing: original draft, Writing: review and editing.

## Declaration of generative AI and AI-assisted technologies in the writing process

During the preparation of this work, the author used Claude Code (Anthropic), an AI-agent software environment powered by the Claude Opus 5 model, for in silico data retrieval, statistical analysis, literature verification and drafting assistance, as described in the Materials and Methods. After using this tool, the author reviewed, independently verified against primary sources, and edited the content as necessary, and takes full responsibility for the content of this publication.

